# Exploring DTI-derived metrics to non-invasively track recellularisation in vascular tissue engineering

**DOI:** 10.1101/2022.07.15.500196

**Authors:** B Tornifoglio, A. J. Stone, P. Mathieu, E. Fitzpatrick, C. Kerskens, C. Lally

## Abstract

Despite significant growth in the field of tissue engineering over the past decades, non-invasive, non-destructive methods to characterise recellularisation of grafts are lacking. Here, we investigate a non-invasive magnetic resonance imaging technique, diffusion tensor imaging (DTI), within acellular and recellularised vascular grafts. Using two decellularised porcine carotid grafts, smooth muscle cells were cultured dynamically for two weeks with terminal time points at day 3, 7, and 14. Grafts were fixed at each time point and investigated by DTI in an *ex vivo* set up. Semi-quantitative histology was carried out to investigate collagen, elastin, and cell density changes over time. DTI-derived metrics, namely the fractional anisotropy, mean diffusivity and tractography, not only were significantly different between day 3 and day 7 grafts, but also distinguished between acellular and recellularised grafts. Specifically, within the wet decellularised grafts, increasing fractional anisotropy was strongly correlated to increasing cell density. The results from this study show, for the first time, DTI’s place in the field of tissue engineering, offering non-invasive, non-destructive insight into graft recellularisation.

## 1.0 Introduction

Interest in tissue engineering small-diameter (<6 mm) vascular grafts has steadily been gaining intense traction over the past 50 years^1–3^. Vascular tissue engineering is believed to be a viable strategy for the treatment of vascular disease in small vessels, for reconstruction or bypass of occlusions and aneurysms or even haemodialysis^1,4,5^. In the absence of favourable autologous grafts^6–8^, which are limited in supply, and in cases where synthetic materials such as Dacron® and polytetrafluoroethylene are not suitable^9,10^, tissue engineered vascular grafts (TEVG) could be the answer. The overarching aim of tissue engineering is to create a 3D scaffold which, once implanted, integrates into native tissue and ultimately becomes indistinguishable in form and function. For small-diameter TEVG this means having similar viscoelastic properties to native arterial tissue, remaining nonthrombogenic, and maintaining adequate patency during its lifetime^2,8,11^.

Both natural^12–15^ and synthetic^14,16–19^ polymers as well as decellularised exogenous grafts^20–28^ have been extensively explored as viable options for small-diameter TEVG. Despite the advances in 3D printing^14^ and electrospinning^16,19^, achieving the native architecture and extracellular matrix of decellularised conduits still remains a challenge^29^. Maintaining native biomechanical properties in arterial grafts means preserving the native collagen and elastin architecture^30^; however, this dense network poses challenges with cell infiltration past a luminal monolayer, both in vitro^25,31,32^ and in vivo^23^. Even when cells were injected directly into this dense extracellular network, cell migration through the media can be very limited^21^. A handful of studies suggest the need for biomechanical preconditioning in vitro in order to aid cell infiltration^16,18,27^; however, infiltration is still limited and seemingly elusive without in vivo studies^20,22,24,28^. With the clear advancement of numerous strategies in vascular tissue engineering, there is a newfound need for versatile, non-invasive, and non-destructive imaging techniques to characterise the TEVG efficacy from in vitro to in vivo.

While each imaging modality offers its own benefits, magnetic resonance imaging (MRI) is one of the few which offers potential for in vitro, preclinical, and clinical applications^33^. MRI offers unparalleled soft-tissue contrast, imaging penetration, safety, and most importantly – can provide anatomical, functional, and cellular information^33–35^. In the last 15 years, a number of studies have been published investigating cell labelling contrasts such as super paramagnetic iron oxide particles^36–41^ or the molecule manganese porphyrin^42,43^. These contrasts have been used with conventional T1-, T2-, and T2*-weighted imaging sequences and all show the potential to identify cell viability as well as the ability to colocalise changes in measured signal with cell migration. Kerans et al. even showed the ability to genetically modify mesenchymal stem cells to produce intracytoplasmic magnetic nanoparticles, a promising methodology which could facilitate MR-based cell tracking^44^. A benchtop-MRI has even been developed which provided detailed insight into diffusion within 3D matrices and inhomogeneities in scaffolds^40^. Multimodal MRI has also been used in cartilage and bone tissue engineering^35,45–48^. Kotecha et al. found that T1 relaxation time and the apparent diffusion coefficient decreased in alginate beads seeded with human chondrocytes over a four week culture, while T2 relaxation times decreased up to week 3 before increasing 8% relative to the initial value^35^. Li et al. demonstrated that quantitative magnetisation transfer imaging metrics, over T1-, T2-, and diffusion weighted imaging metrics, correlated with increased glycosaminoglycans in gelatin sponges seeded with mesenchymal stem cells. Cheng et al. similarly used T1-, T2-, and diffusion weighted imaging to successfully differentiate between acellular matrices with and without hyaluronic acid^49^. Studies have also used spatial variation in MRI contrasts to investigate how well constructs integrate into host tissue^46,47,50^ for cartilage and bone regeneration. To the author’s knowledge, MR diffusion tensor imaging (DTI) has only been applied to tissue engineering studies once^17^. Ghazanfari et al. used DTI, validated by confocal laser scanning microscopy, to monitor the evolution and reorientation of collagen fibres on polyglycolic acid meshes of different aspect ratios seeded with human vascular derived cells for up to six weeks^17^. While not specific to tissue engineering, Gimenez et al. used both in vivo and ex vivo DTI to detect glioma cell migration in mice brains^51^, highlighting a sensitivity to the presence of cell content and migration.

DTI is an advantageous MR technique capable of probing tissue microstructure and is sensitive to changes in the underlying microstructure. The aim of this study was to evaluate recellularisation in vascular grafts, specifically decellularised porcine carotids, over a two-week culture period using just one MRI technique - DTI. Having previously been shown to be sensitive to specific changes in arterial microstructure^52,53^, the DTI results presented in this work are explained and validated by quantitative histological analysis. This work shows that both qualitative and quantitative DTI-derived metrics are not only sensitive to changes in microstructure in TEVG, but strongly correlate with the recellularisation of these grafts. The novel findings in this study establish the exciting and transformative potential of DTI within the field of tissue engineering.

## 2.0 Methods

A brief overview of the methods in this study are presented in Figure 1.

**Figure 1.**
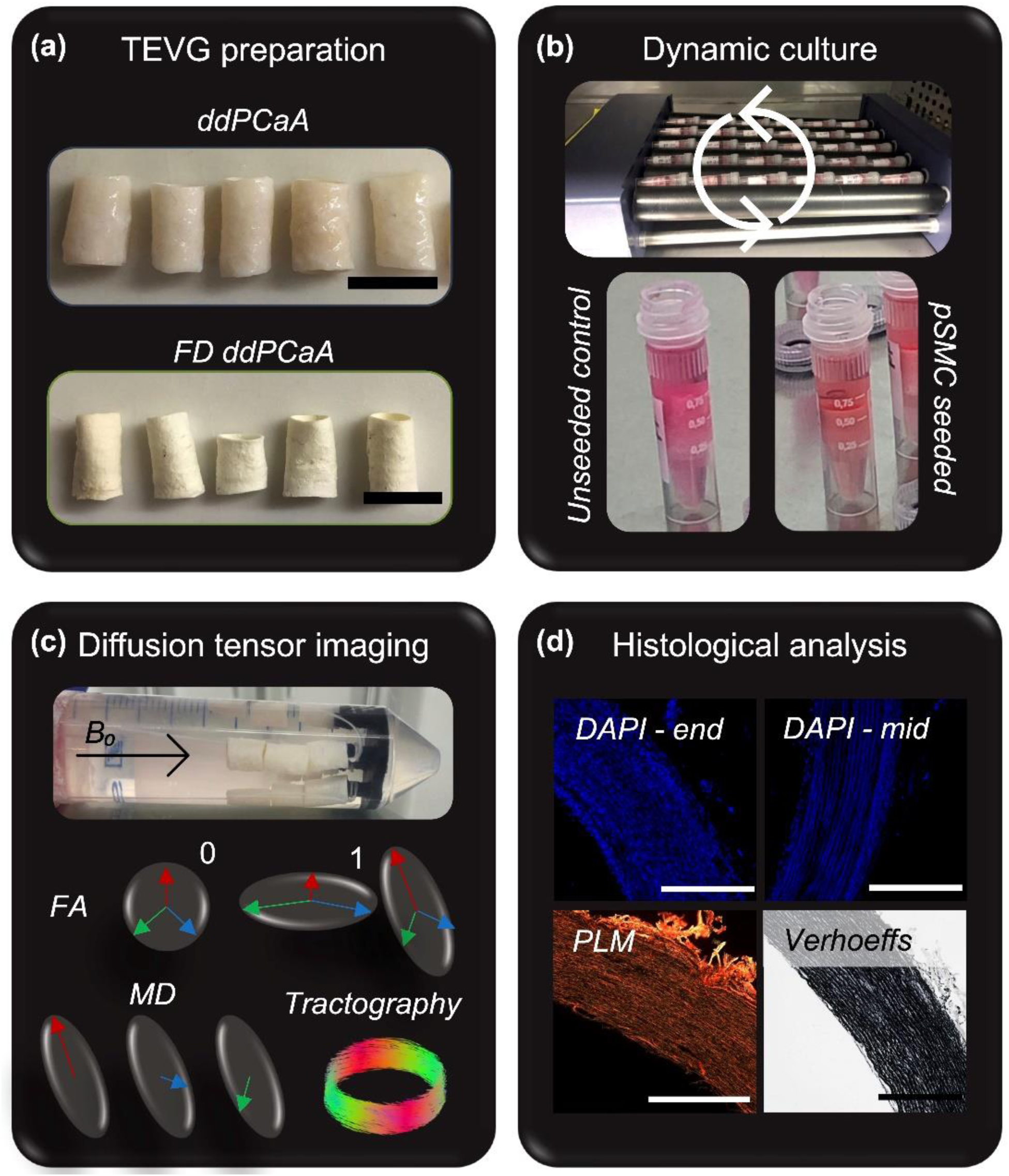
General overview of the presented study. (a) Two TEVGs were used in this study: decellularised, de-intima porcine carotid arteries (ddPCaA) and freeze-dried ddPCaA (FD ddPCaA). Scale bars 10 mm. (b) TEVGs were seeded with pSMC and cultured in individual eppendorf tubes on a roller for up to two weeks. (c) Diffusion tensor imaging (DTI) was performed on both TEGV groups to calculate the DTI-derived fractional anisotropy (FA) and mean diffusivity (MD) as well as to perform tractography. (d) Qualitative and quantitative histological analysis was performed on each graft. Scale bars are 500 μm.

### 2.1 TEVG preparation

Fresh porcine carotids were excised from 6-month-old healthy Large White pigs from the same abattoir, cleaned of connective tissue, and cryopreserved within 3 hours of sacrifice. Tissue freezing medium was used in cryopreservation in order to prevent the formation of ice crystals and maintain native tissue microstructure^52,54^. Vessels were thawed at 37°C and, using previously reported methods, decellularised^32,52,55^. Briefly, vessels were perfused with 0.1 M sodium hydroxide at 2 Hz for 15 hours via a peristaltic pump followed by 0.1 M sodium chloride for 32 hours. Perfusion was done at a pressure of 100 mmHg maintained by a water column. Vessels were then treated with 10 μl/ml DNAase (LS006342, Worthington Biochemical Corporation) and 2 μl/ml primocin (Ant-pm-2, InvivoGen) at 37°C for 19 hours, followed by a phosphate buffered saline (PBS) wash. Decellularised vessels were cut into 10 mm long segments and inverted to remove the intimal layer. This exposed the highly aligned microstructure of the medial layer of the arterial wall^56–58^. A scalpel was carefully dragged along the intima layer, with no pressure applied, and fine tipped forceps were used to peel the intimal layer off. Any vessels which had uneven intimal layer removal were excluded. Vessels up to this point constitute one group used in this study – wet decellularised, de-intima porcine carotid arteries (ddPCaA), seen in Figure 1(a). Other wet vessels, following the same decellularisation preparation, were lyophilized using a similar protocol to one previously reported which preserves the mechanical properties of the scaffold^59^. Briefly, vessels were allowed to airdry then snap-frozen in liquid nitrogen and transferred immediately into a precooled freeze drier (FreeZone, Labconco Corporation, Kansas City, MO) at -40°C. The samples were kept at -40°C for two hours and then ice crystal sublimation was induced by decreasing the pressure to 0.266 mbar and increasing the temperature to 0°C at 1°C/min. After 18 hours, a secondary drying phase to 20°C at 1°C/min concluded the lyophilization process. These scaffolds constitute the second graft group in this study – freeze-dried, decellularised, de-intima porcine carotid arteries (FD ddPCaA), seen in Figure 1(a). All vessels were sterilised in 100% ethanol for one hour, rinsed in sterile PBS and then transferred into sterile culture medium and incubated statically at 37°C for 24 hours prior to seeding.

### 2.2 Cell seeding and dynamic culture

Porcine smooth muscle cells (pSMC) were isolated using a similar protocol used previously for rat smooth muscle cells^60^. Porcine carotid arteries were obtained from the abattoir, cleaned of connective tissue, and the adventitia was removed. Carotids were cut into segments no larger than 1 mm^3^ and placed into MgCl_2_ + CaCl_2_ supplemented PBS with 0.7 mg/ml collagenase type 1A from Clostridium histolyticum (Sigma) and 0.25 mg/ml elastase type III from porcine pancreas (Sigma). The tissue was digested under constant agitation at 37°C until totally digested, approximately 7 hours. The resulting cell suspension was spun down at 400 g and the digestion solution removed. The pSMC were cultured in high glucose DMEM with Glutamax (Biosciences) with 10% FBS (Gibco) and 0.2% primocin at 37°C, 5% CO_2_, 20% O_2_ and used at passage 6 for all experiments. For TEVG seeding, a 1 ml eppendorf was used for individual vessels, which were seeded at a concentration of 1.5·10^6^ cells/ml. Each eppendorf was fully closed and placed on a roller in an incubator at 37°C (Figure 1(b)). Due to the closed nature of the culture, media was changed daily in the A.M. and the vessels were vented in the P.M., approximately 10 hours later, for 5-10 minutes in sterile conditions. The dynamic culture lasted two weeks with terminal time points on day 3, 7, and 14. There were four vessels per time point for both ddPCaA and FD ddPCaA and one unseeded control for both groups at each time point. Vessels were fixed in 10% formalin for three hours at their respective time points then transferred to PBS until imaging.

### 2.3 DTI acquisition and analysis

All MR imaging was performed with a small bore horizontal 7 Tesla Bruker BioSpec 70/30 USR system (Bruker, Ettlinger Germany) equipped with a receive only 8-channel surface array coil, birdcage design transmit coil, shielded gradients (maximum strength 770 mT/m) and Paravision 7 software. Dental floss was tied onto a custom-made 3D printed holder which allowed for 12 vessels – three time points (n=3 per time point) and one control per time point – to be imaged together (Figure 1(c)). This set up was secured into a 50 ml falcon tube with fresh PBS for imaging. All ddPCaA vessels were imaged in one scan session and similarly all FD ddPCaA vessels were imaged together: two scan sessions total. A standard 3D spinecho DTI sequence with monopolar gradients was used with the following parameters: TE/TR: 17.682/1000 ms, averages: 1, image size: 96 × 96 × 96, field of view 30 × 30 × 30 mm, resolution: 0.313 × 0.313 × 0.313 mm, b values: 0, 800 s/mm^2^, 10 isotopically distributed directions and a total acquisition time of 28 hours and 9 minutes. Imaging was performed at room temperature, approximately 25°C.

All raw data was processed as previously described^52^. Fractional anisotropy (FA) and mean diffusivity (MD) were calculated in ExploreDTI^61^. Due to variations in sample lengths and regions outside the imaging field of view, all vessels were analysed for the same number of slices (eight slices – 2.5 mm of length); these slices started at one end of the vessel and moved inwards longitudinally towards the centre of the vessel. FA and MD were averaged across the slices for each vessel, yielding one mean and standard deviation per vessel. Tractography was performed in ExploreDTI and was done on the same eight slices. The tracking parameters were as follows: seed point resolution: 0.3125 × 0.3125 × 0.3125, seed FA threshold: 0.1; FA tracking threshold range: 0.1-1, MD tracking threshold: 0-infinity, linear, planar, and spherical geometry tracking threshold range: 0-1, fibre length range: 0.5-50 mm, angle threshold: 45°, step size: 0.3125, linear interpolation method, and no random perturbation of seed points. Representative slices are presented in parametric maps (FA and MD) and representative vessels shown for tractography. Slice to slice variations are presented in Supplementary figure 1.

### 2.4 Microstructural quantification

One vessel at each time point was qualitatively imaged with DAPI (Sigma) staining prior to MR imaging on a Leica SP8 scanning confocal microscope. TEVG were sectioned into 2 mm long segments and if possible, images were taken both at the end of the vessel (exposed edge) and the middle of the vessel. All images were taken at an image size of 1024 × 1024 and a scan speed of 400 Hz. These images were used to check for successful cell seeding and monitor cell infiltration during culture; these vessels were not imaged with MRI or used for quantitative histology. They are qualitative 3D representations of recellularisation for each time point.

Quantitative histological analysis was performed after MR imaging for all vessels. Grafts were brought to histology and sectioned at 8 μm slices prior to DAPI, picrosirius red, and Verhoeff’s elastin staining. Brightfield imaging of Verhoeff’s elastin was done on an Aperio CS2 microscope with ImageScope software V12.3 (Leica Biosystems Imaging, Inc., Vista, California). Elastin content (specifically the stained area fraction of the histological sections) was determined using QuPath and a previously established method^62^; specifically using very high (0.99 μm/pixel) resolution, the Verhoeff channel (determined by the stain vector) and a threshold of 1. Elastin content was determined across three regions of interest per vessel and averaged to yield one value per vessel. Polarised light microscopy (PLM) was performed on picrosirius red stained samples to maximise the birefringence of collagen (exposure time 1100 ms) using an Olympus BX41 microscope with Ocular V2.0 software (Teledyne Photometrics, Tucson, Arizona) and used to determine collagen content using a previously established method^63^. Three regions of interest were used per vessel for collagen content determination. DAPI stained histology slides were imaged on an Olympus BX51 upright microscope (exposure time 200 ms). Cell counting was done manually using the cell counter plug-in in ImageJ^64^ and tissue area measured using tissue detection in QuPath^65^. Cell densities were calculated across nine regions of interest per vessel and averaged to yield one density per vessel. Alcian blue staining was also performed to qualitatively look at the GAG content in the control grafts and are presented in Supplementary figure 2.

**Figure 2.**
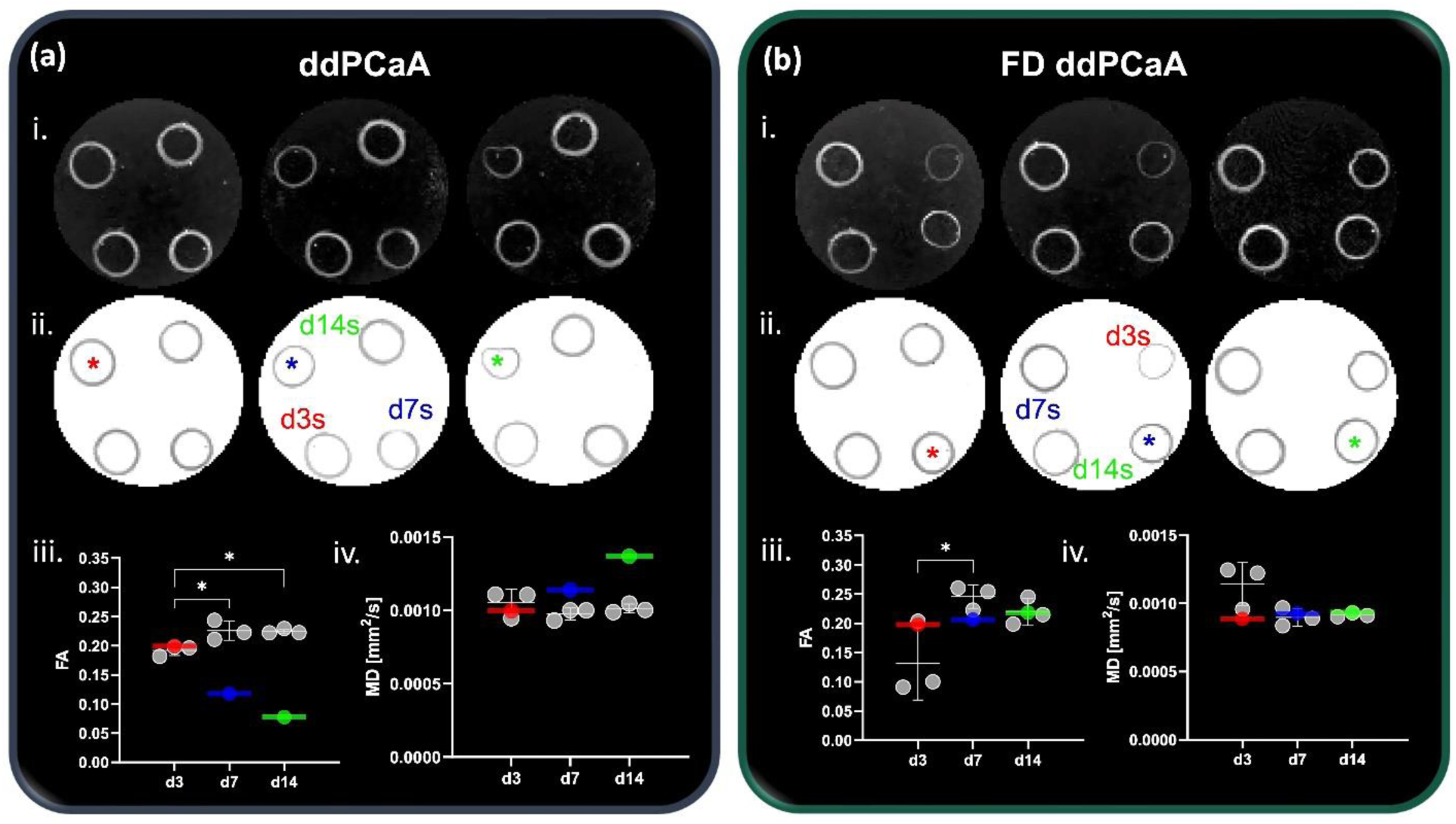
DTI-derived metrics for (a) ddPCaA and (b) FD ddPCaA. (a, b-i) FA and (a, b-ii) MD parametric maps and mean (a, b-iii) FA and (a, b-iv) MD metrics for each time point. Windowing of parametric maps matches the axes in graphs. Coloured asterisks (in parametric maps) and lines (in graphs) represent unseeded controls for each time point (n=1). Red signifies day 3 control, blue is day 7, and green day 14. Significance between TEVG (n=3 per time point) was determined with ordinary one-way ANOVAs with Tukey’s multiple comparisons. (a-iii) *p=0.0216 between d3 and d7 and *p=0.0248 between d3 and 14. (b-iii) *p=0.0308.

### 2.5 Statistical analysis

All statistical analysis was performed using GraphPad Prism V9.3.1. Normality was tested using D’Agostino-Pearson tests (alpha=0.5) and equal variances using Brown-Forsythe tests. FA, MD and tractography measures all passed both normality and equal variances so ordinary one-way ANOVAs with Tukey’s post hoc multiple comparisons were used. All DTI-derived data is presented as mean per vessel (averaged over eight slices) and standard deviation across the three vessels per time point. Similarly, quantitative histology measures were tested for normality and equal variances; all data passed both tests and ordinary one-way ANOVAs with Tukey’s post hoc multiple comparisons tests were used. To investigate the relationship between FA and MD with cell density and collagen-to-elastin ratio, Pearson’s correlations were tested, where an r value greater than 0.7 was considered a strong correlation^66^. The ROUT method (Q=1) was used to check for outliers, from which none were found^67^.

## 3.0 Results

### 3.1 DTI-derived metrics of recellularised porcine carotid arteries

In this study, two different TEVGs, ddPCaA and FD ddPCaA, were tested to facilitate recellularisation and explored via DTI to non-invasively track cell infiltration and growth during a two-week culture period.

Figure 2 presents DTI-derived FA and MD parametric maps and mean values for each time point within the two groups. Panel (a) shows the results for ddPCaA. Visually, both parametric maps, (a-i) FA and (b-ii) MD, show qualitative differences between unseeded controls (top left within each slice, marked with asterisks) and different time points in the two-week culture. These differences were quantified and are presented in (a-iii) and (a-iv). The mean FA of vessels cultured for 3 days was 0.19 ± 0.0087, which was significantly lower than that of day 7 vessels, 0.23 ± 0.016, and day 14 vessels, 0.22 ± 0.0041. Interestingly, the mean FA of the unseeded controls consistently decreased between time points. The day 3 unseeded control had an FA of 0.19, the day 7 – 0.12, and the day 14 vessel – 0.08 (visualised by the coloured lines in (a-iii)). The opposite trend was seen with MD for the different time points. Day 3 seeded vessels had the highest MD, 0.0011 ± 0.00009 mm^2^/s, followed by day 14 vessels, 0.0010 ± 0.00004 mm^2^/s, and day 7, 0.00098 ± 0.00003 mm^2^/s. Increasing MD was also observed with the unseeded controls from day 3 to 14, 0.00099, 0.0011, and 0.0014 mm^2^/s, respectively.

The same metrics for the FD ddPCaA are presented in Figure 2 panel (b). Parametric maps of FA and MD are shown in (b-i) and (b-ii). When quantified, the FA of day 7 vessels was significantly higher than day 3 vessels, 0.25 ± 0.02 and 0.14 ± 0.063, respectively. No significant difference was seen when compared to day 14 vessels, 0.22 ± 0.023. Unlike ddPCaA, the FA of the unseeded FD ddPCaA controls remained constant from day 3 to 14: 0.19, 0.21, 0.22, respectively. No significant differences were seen from day 3 to 14 with respect to MD of the pSMC seeded FD ddPCaA, 0.0011 ± 0.00016, 0.00089 ± 0.00007, and 0.00092 ± 0.00001 mm^2^/s, respectively. Again, the unseeded controls remained constant from day 3 to 14: 0.00089, 0.00093, and 0.00093 mm^2^/s, respectively.

Tractography was performed for all TEVGs and results for unseeded controls as well as representative grafts for each time point are presented in Figure 3. In both panel (a) and (b) the top row presents the unseeded controls while the second row is pSMC seeded grafts. Qualitatively, for both groups there is a lack of continuity between tracks in the controls that is not as apparent in the recellularised vessels. Differences between time points was quantified in (a, b-ii-iv). The coloured lines in (ii-iv) are the unseeded controls which, with the exception of the day 3 control, are all well-below the means for the pSMC seeded vessels at day 7 and 14 for ddPCaA Figure 3((a)ii-iv). Day 7 pSMC seeded ddPCaA had a significantly higher (a-ii) tract volume and (a-iii) number of tracts than day 3 vessels. Day 14 ddPCaA also had a significantly higher (a-iii) number of tracts than day 3 vessels. Similarly, day 7 FD ddPCaA had significantly higher (b-ii) tract volume and (b-iii) number of tracts than day 3 vessels. Day 14 FD ddPCaA vessels also had significantly higher (b-ii) tract volume and (b-iii) number of tracts than day 3 vessels.

**Figure 3.**
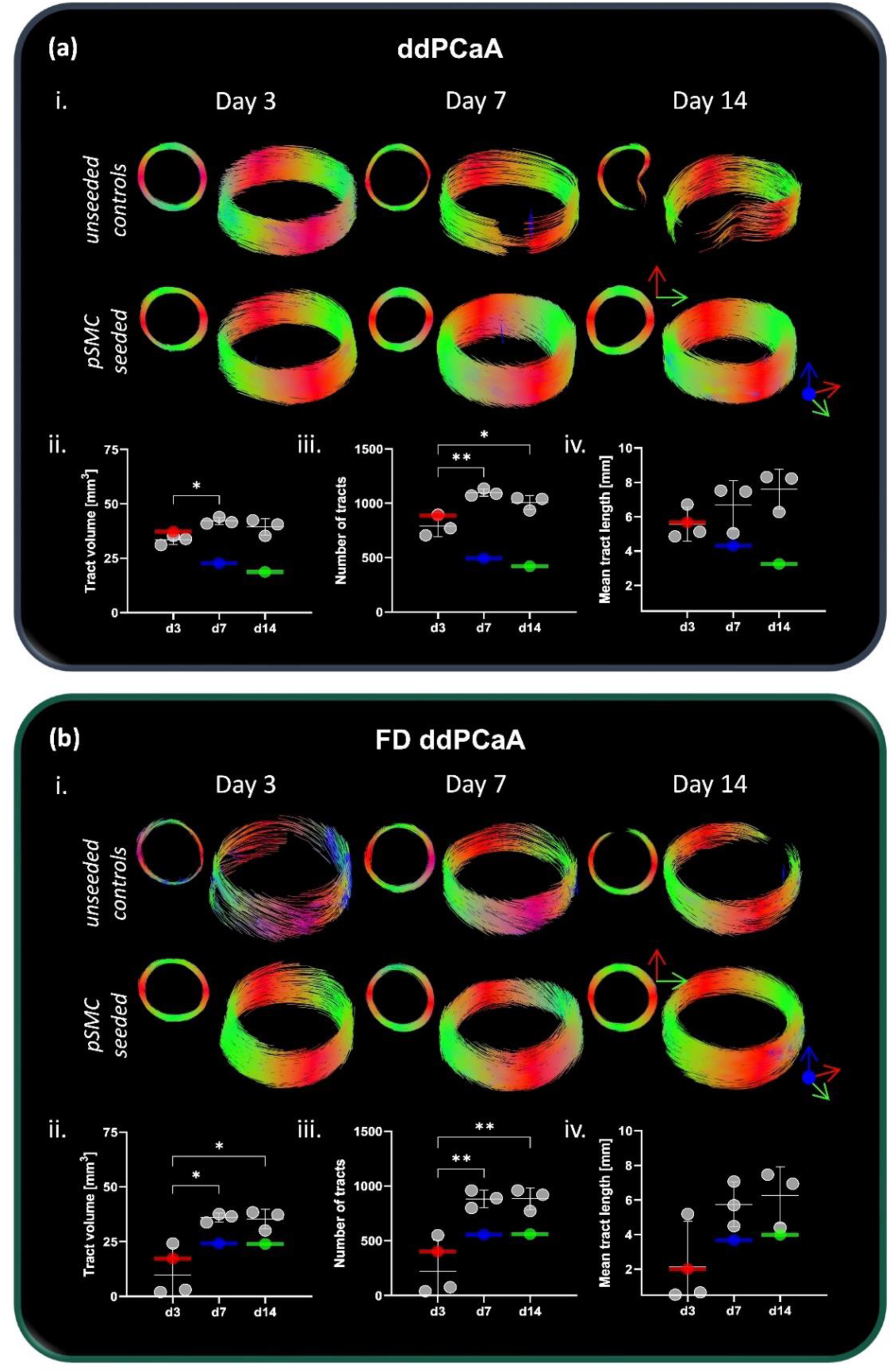
Tractography for both (a) ddPCaA and (b) FD ddPCaA TEVGs. (a, b-i) Unseeded control vessels are shown in the top rows, with representative recellularised grafts in the second row for each group. (a, b-ii) Mean tract volume, (a, b-iii) number the tracts, and (a, b-iv) mean tract length for both groups. Significance was determined with ordinary one-way ANOVAs with Tukey’s post hoc multiple comparisons: (a-ii) *p=0.0160, (a-iii) *p=0.0204, **p=0.0039, (b-ii) *p=0.0144 between d3 and d7, *p=0.0163 between d3 and d14, (b-iii) **p=0.0100 between d3 and d7, **p=0.0099 between d3 and d14. Coloured lined are unseeded controls for each time point (n=1).

### 3.2 Qualitative histology

At day 3, 7, and 14 one TEVG from each group was DAPI stained and imaged for a qualitative look at cell infiltration. Figure 4 highlights these findings. For both groups through the thickness cell infiltration was seen on day 3. Thorough infiltration through the wall thickness was seen in Figure 4(a) day 7 and 14 ddPCaA grafts at the exposed edges; however, when looking from a more central region of the graft, cells were only present on the luminal and adventitial sides. Similar results were seen for Figure 4(b) FD ddPCaA; however, the day 14 graft showed no infiltration through the wall thickness. Additionally, the luminal cell layer was thicker than a monolayer for both day 7 and 14. Alcian blue staining of unseeded controls showed overall very faint GAG staining, but a slight decrease in the already low GAG content of the control grafts along the lumen (Supplementary figure 2). The ddPCaA grafts appear to have increased intra-wall separation when compared to the FD ddPCaA grafts, noted by an asterisk. The FD ddPCaA grafts appear more densely compacted within the vessel wall.

**Figure 4.**
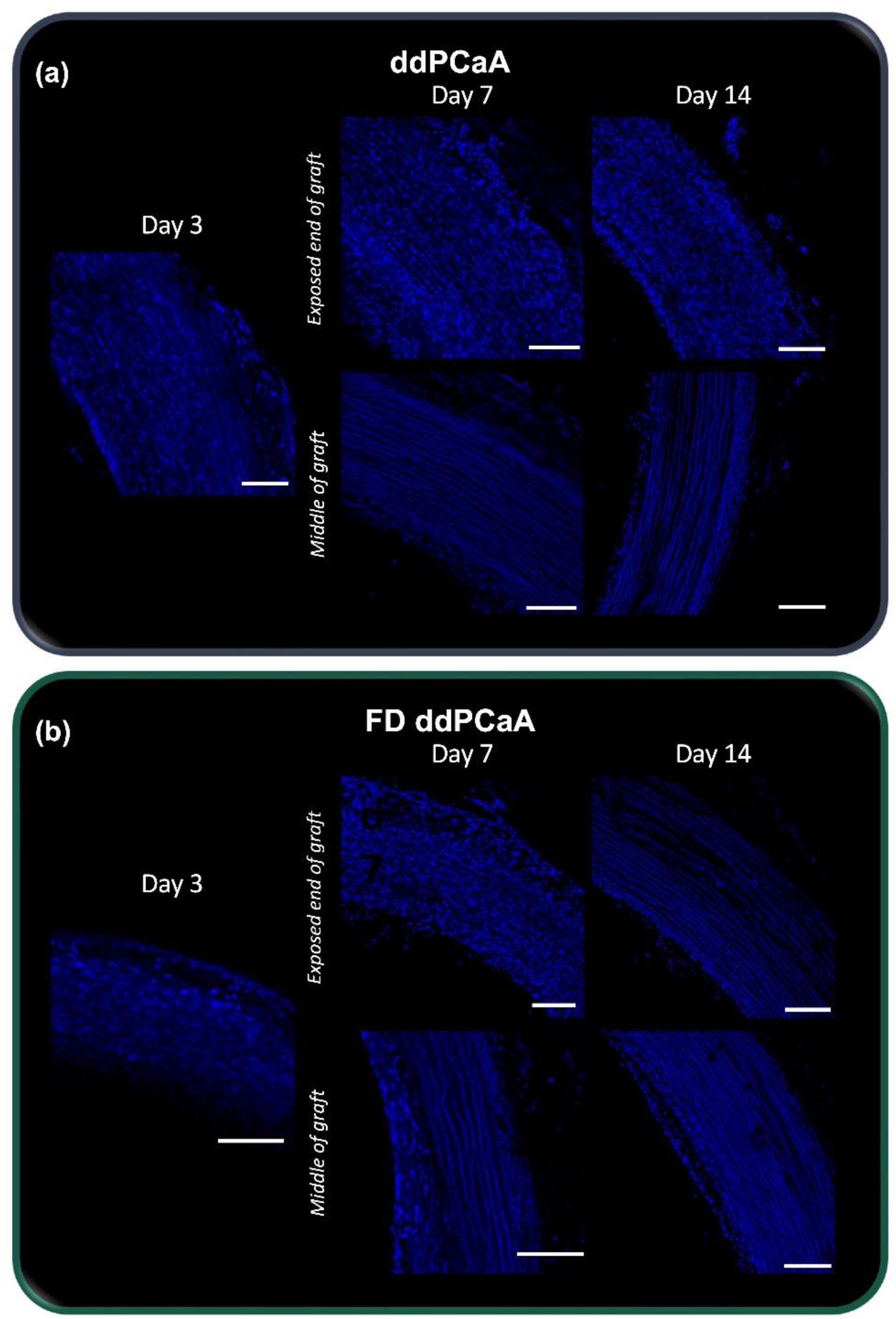
Qualitative scanning confocal microscopy of DAPI stained TEVG at each time point. (a) ddPCaA and (b) FD ddPCaA both showed significant cell infiltration up to a depth of 400 μm at the exposed ends of the grafts (top rows for day 7 and 14). However, when looking at edges from the centres of the grafts, this infiltration was less apparent, and a luminal layer of cells was present (bottom row for day 7 and day 14). Scale bars are 200 μm.

### 3.3 Quantitative histology

Representative histology is shown for ddPCaA in Figure 5Figure and for FD ddPCaA in Figure *6*6. While no significant difference was found, increasing cell density was observed for ddPCaA (Figure 5(a)). The collagen-to-elastin ratio remained constant during culture for ddPCaA, also with no significant differences. Due to visibly inconsistent cell seeding observed in the FD ddPCaA, the top row of Figure *6*6 has both unseeded controls in the top right subfigure and examples of localised high cell density regions in the bottom subfigure. Similar to ddPCaA, no significant differences were found for FD ddPCaA (Figure *6*6(a, b)), yet the cell density and collagen-to-elastin ratio both dipped down at day 7 before increasing again at day 14 but this was not significant.

**Figure 5.**
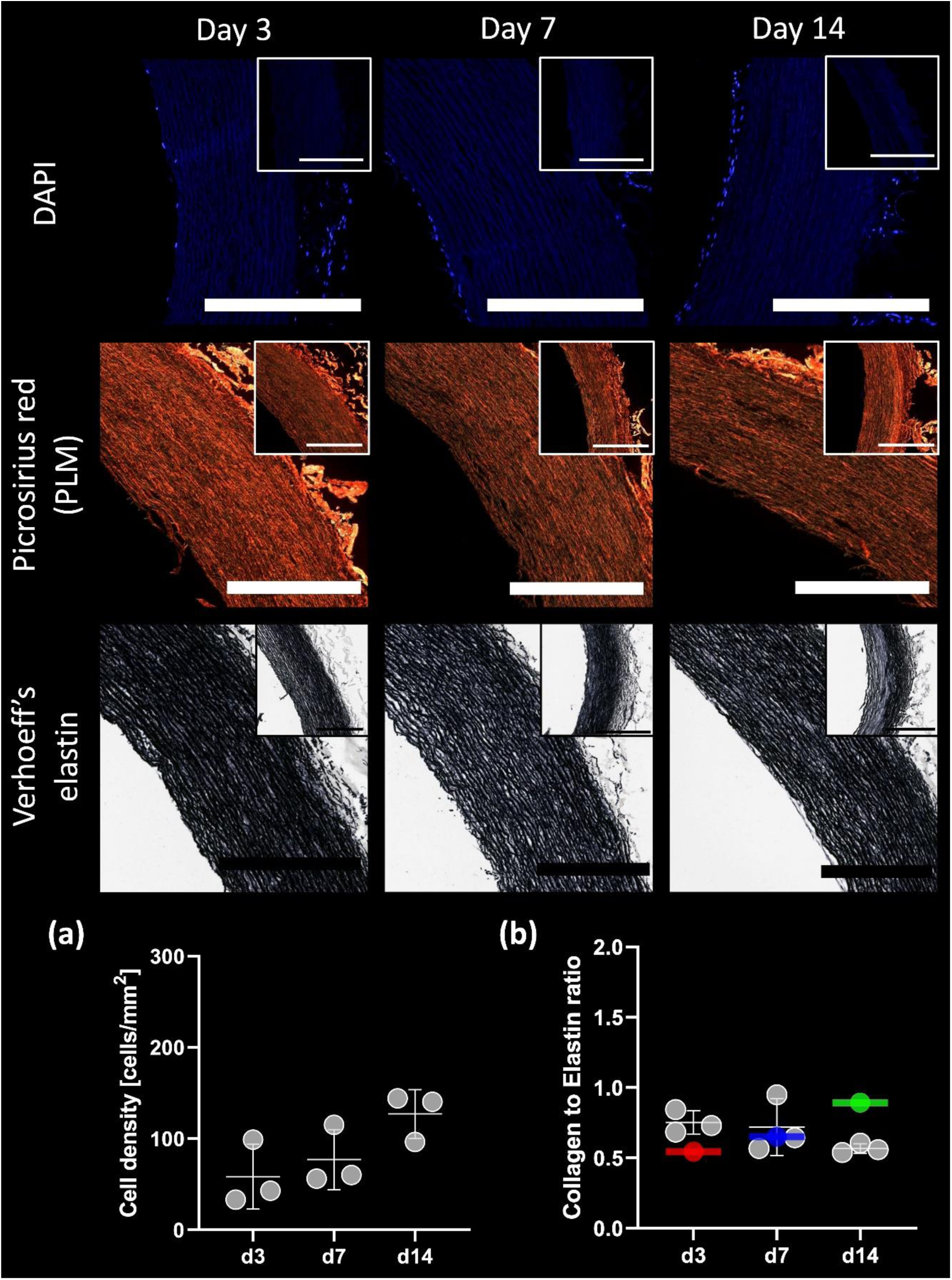
Representative histology of ddPCaA. DAPI stained cross-sections (top row), picrosirius red stained sections imaged with PLM (second row) and Verhoeff’s-stained cross-sections (third row) at each time point. Subfigures are unseeded controls at each time point. All scale bars are 500 μm. (a) Cell densities (b) and collagen-to-elastin ratios for all ddPCaA TEVG. Coloured lines are unseeded controls. No significant differences were found using ordinary one-way ANOVAs with Tukey’s multiple comparisons.

**Figure 6.**
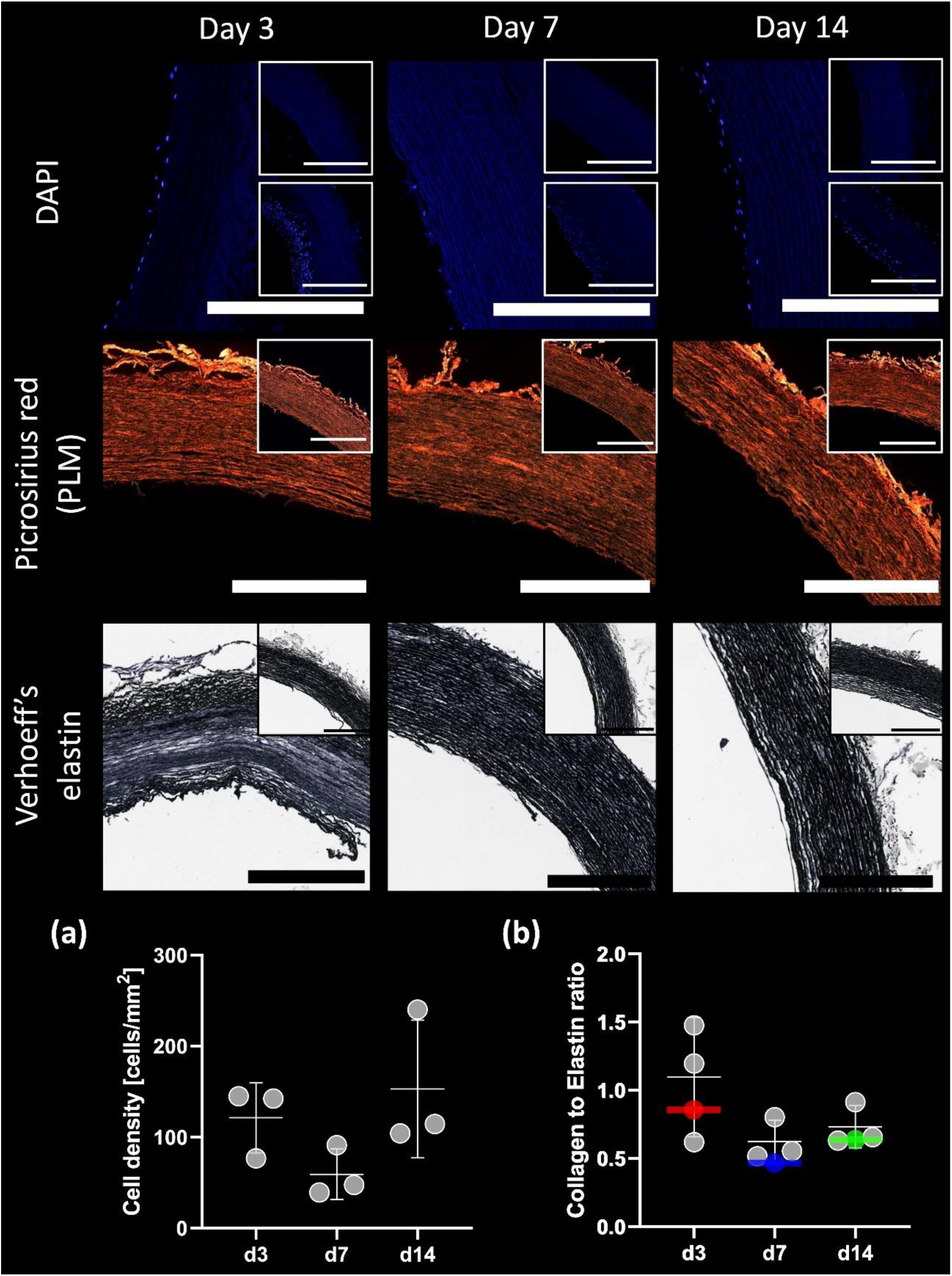
Representative histology of FD ddPCaA. DAPI stained cross-sections (top row), picrosirius red stained sections imaged with PLM (second row) and Verhoeff’s-stained cross-sections (third row) at each time point. Subfigures are unseeded controls at each time point. DAPI subfigures are unseeded control (top) and examples of densely seeded localised regions in recellularised grafts (bottom). All scale bars are 500 μm. (a) Cell densities (b) and collagen-to-elastin ratios for all ddPCaA TEVG. Coloured lines are unseeded controls. No significant differences were found using ordinary one-way ANOVAs with Tukey’s multiple comparisons.

Figure 7 presents the correlations between DTI-derived FA and MD with quantitative histology measures. Panel (a) shows ddPCaA and FD ddPCaA is shown in panel (b). For ddPCaA grafts, cell density was strongly and significantly correlated to FA (Figure 7(a)-i). For the FD ddPCaA the collagen-to-elastin ratio was strongly correlated with both FA and MD (Figure 7(b)-i, ii).

**Figure 7.**
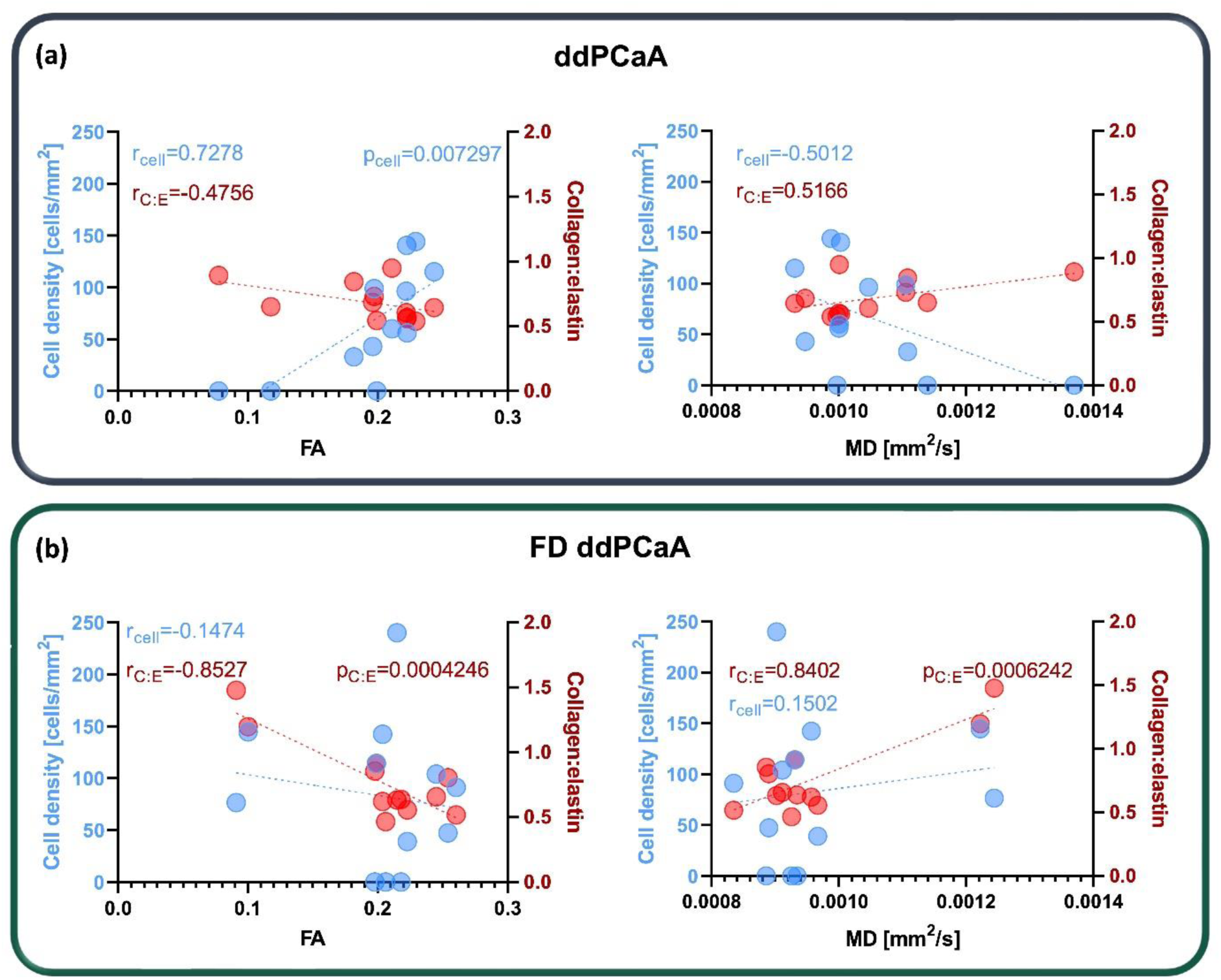
DTI-derived FA and MD correlations with quantitative histology. (a) FA and MD correlations with cell density and collagen-to-elastin for ddPCaA and (b) for FD ddPCaA. Pearson’s r values are presented for strong correlations alongside significant p values. Each data point corresponds to the average DTI-derived metric and quantitative histological measure per vessel (n=3 per time point, n=1 control per time point).

## 4.0 Discussion and conclusion

Non-invasive and non-destructive characterisation within tissue engineering is not only an advantageous goal, but also an attainable one. The ability to track recellularisation – both in quantity of cell repopulation and in quality of recellularised microstructure – would allow for non-destructive, non-terminal time points which can dictate experimental planning. The potential benefits, while significant in the transition from in vitro to preclinical studies, are also relevant in the clinical translation. It would be transformative to be able to use one consistent, reliable characterisation technique during translation, and this could also positively address the 3Rs^68^. MRI offers a safe and non-ionizing imaging method which has the potential to impact preclinical experimental design by reducing the sample numbers needed by eliminating early terminal time points.

To date, the biomedical field has very few examples of MRI for tissue engineering applications. When it comes to solely using the intrinsic contrast provided by the abundance of ^1^H protons, the majority of studies use multiple contrasts^35,45,48,49^ and it can be difficult to elucidate what changes in the tissues are resulting in the consequent changes in measurable signal. Chemical exchange saturation transfer has shown an ability to measure specific concentrations of different molecules^69,70^; however, its use in tissue engineering is limited^71^. DTI has been used in a handful of focused studies in different tissues to understand how different microstructural components affect the measurable signal^52,53,72^. A significant benefit to DTI is the ability to get directional information about the underlying microstructure – a feature which is highly relevant in both cardiac^19,73–75^ and cartilaginous^76–78^ tissues.

The results in this study showed, for the first time, that DTI-metrics are capable of tracking recellularisation in vascular grafts. Specifically, for ddPCaA the FA was significantly higher in day 7 and 14 grafts compared to day 3 grafts. Additionally, for day 7 and 14 grafts, the FA was considerably lower in the unseeded controls – pointing to the ability to differentiate between recellularised and acellular grafts. As collagen and elastin content did not change between time points, the increasing cell density, while not significant, was the dominant microstructural change. This relationship is further confirmed by the strong correlation between FA and cell density. While the mono-exponential DTI model has shown that the main contributor to the measurable anisotropic diffusion in arterial tissue is cellular content^52^, non-gaussian schemes, such as the stretched-exponential model, have been used to further hone in on alterations in the tissue microstructure^53^.

Only one study has looked at the relationship with cell density related to tumour progression and DTI-derived perpendicular diffusivity. Those authors identified both perpendicular diffusivity and FA as potential markers for glioma cell migration and invasion^51^. Glioma cells migrate along white fibre bundles in the brain, and when these are disrupted due to erosion, destruction, impingement, and peritumoral edema in tumour progression, the disordered microstructure is identifiable by decreased FA^79,80^.

Interestingly, for the FD ddPCaA, while the FA was significantly higher in day seven grafts, there was a drop in cell density and collagen, and increase in elastin compared to day 3 grafts. Previous work has shown a strong correlation between FA and elastin which likely explains this finding^62^. With the layered structure and highly aligned collagen and elastin present within the arterial wall, the sublimation of ice crystals does not necessarily yield pore formation from lyophilisation^59^, as typically seen in homogenous solutions, and no disruption to this microstructure was seen histologically. While lyophilisation is typically used to achieve high levels of porosity and anisotropy^74^, the protocol used in this study instead focused on maintaining mechanical integrity of the graft rather than creating larger pores^59^. Further studies should be done to investigate varying porosities in TEVGs and their influence on DTI-derived parameters.

It is interesting to note the decreasing FA and increasing MD in ddPCaA unseeded controls alongside less coherent tractography. Over a two-week period in culture medium, at 37°C, and on a dynamic roller, it would not be surprising to have tissue degradation. However, no clear degradation was observed regarding collagen or elastin degradation in the unseeded controls. While GAG content was not quantitatively investigated, unseeded controls qualitatively appeared to have less GAG from day 3 to day 14 which could explain this result (Supplementary figure 2). Upon visual inspection of tractography of the unseeded control vessels between the two groups it becomes clear that the overall quality decreases in ddPCaA while no clear trend is seen in the FD ddPCaA controls. When looking between the two different graft types, the FD ddPCaA appear more densely packed within the wall, seen in supplementary figure 2, compared to the ddPCaA grafts. It is possible the lyophilisation compacts the wall, leading to a denser tissue which protects the graft from degradation as seen in the DTI-metrics for the ddPCaA grafts (Figure 2(a) and Figure 3(a)). Unquestionably, however, the tractography of the recellularised grafts for both groups at all time points is more representative of native arterial tissue^52^ than the decellularised control TEVGs. These differences were further shown in the quantification of tractography based measures. Tractography has previously been shown to be a useful visualisation tool when it comes to crude^81^ and specific^52^ microstructural changes in healthy and diseased^82^ arterial tissue. While it is clinically used in the brain^79^, DTI tractography has demonstrated sensitivity to pathological processes in the spinal cord^83^ and results from this study further its potential use to the field of tissue engineering.

The samples in this study were fixed at each time point due to the lengthy scan time at ambient temperature, in a non-sterile environment – all limiting factors for continuing a cell culture experiment. Future work aims to develop an MRI compatible bioreactor which would allow for TEVG to be taken out of culture, imaged, and returned to culture. Novel MRI-compatible bioreactors already exist^84^ and with the wide-spread use of 3D printing and MRI-compatible sensors^85^, there are exciting opportunities to harness MRI as a longitudinal imaging technique in tissue engineering. It has been shown how vascular stem cells align with underlying collagen fibres regardless of the direction of strain imposed; however, when this alignment is parallel to the direction of strain – it has been shown to result in increased proliferation^86^. Ghazanfari et al. also found that higher aspect ratio constructs resulted in more highly aligned collagen, and likely cells. While the current study only used circumferential rotation during culture, and it proved sufficient to facilitate cellular alignment with the TEVG microstructure, future studies should investigate more physiological mechanical loading conditions. DTI is notoriously tricky to implement in vivo outside of the brain due to physiological motion and lengthy scan times required to obtain high angular resolution data, making static tissue engineering applications the ideal candidate for this imaging technique in the interim.

The results presented in this study clearly demonstrate the use of DTI-derived metrics in the field of tissue engineering. We show that both FA and tractography measures are sensitive to recellularisation of decellularised TEVG, both qualitatively and quantitatively. Not only can DTI metrics track recellularisation longitudinally, but these metrics can differentiate between acellular and recellularised grafts. Findings from this study establish the feasibility of DTI to be used in tissue engineering and highlights the exciting potential of this technique for accurate non-invasive and non-destructive characterisation within TEVGs.

## Author contributions

B.T. performed all experimental work and data analysis. P.M. isolated the cells used in this study. B.T., A.J.S. and C.K. contributed to the development of the DTI protocol. C.L. conceived and supervised the study while C.L. and B.T. contributed to the study design. All authors reviewed the manuscript.

