## Supplementary material for "Exploring DTI-derived metrics to non-invasively track recellularisation in vascular tissue engineering"

**Affiliations:**
[1]Trinity Centre for Biomedical Engineering, Trinity Biomedical Sciences Institute, Trinity College Dublin, Ireland; [2]Department of Mechanical, Manufacturing and Biomedical Engineering, School of Engineering, Trinity College Dublin, Ireland; [3]Department of Medical Physics and Clinical Engineering, St. Vincent's University Hospital, Dublin, Ireland; [4]Trinity College Institute of Neuroscience, Trinity College Dublin, Ireland; [5]Advanced Materials and Bioengineering Research Centre (AMBER), Royal College of Surgeons in Ireland and Trinity College Dublin, Ireland

### Supplementary material

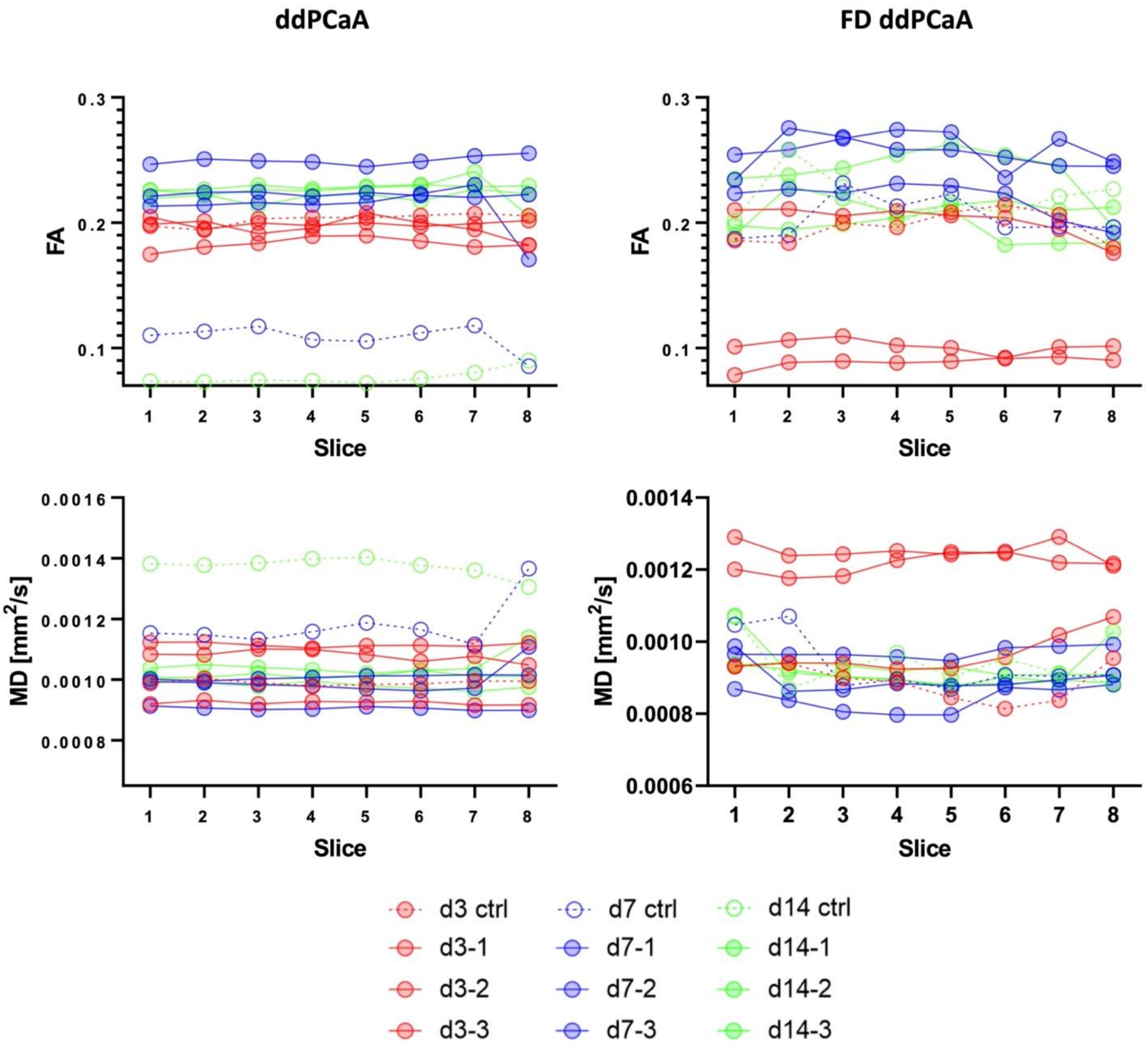


Supplementary figure 1. Mean FA and MD values per slice for each vessel. Red are day 3 vessels, blue day 7, and green day 14. Dotted lines connect the mean values for the unseeded controls while solid lines connect recellularised grafts.

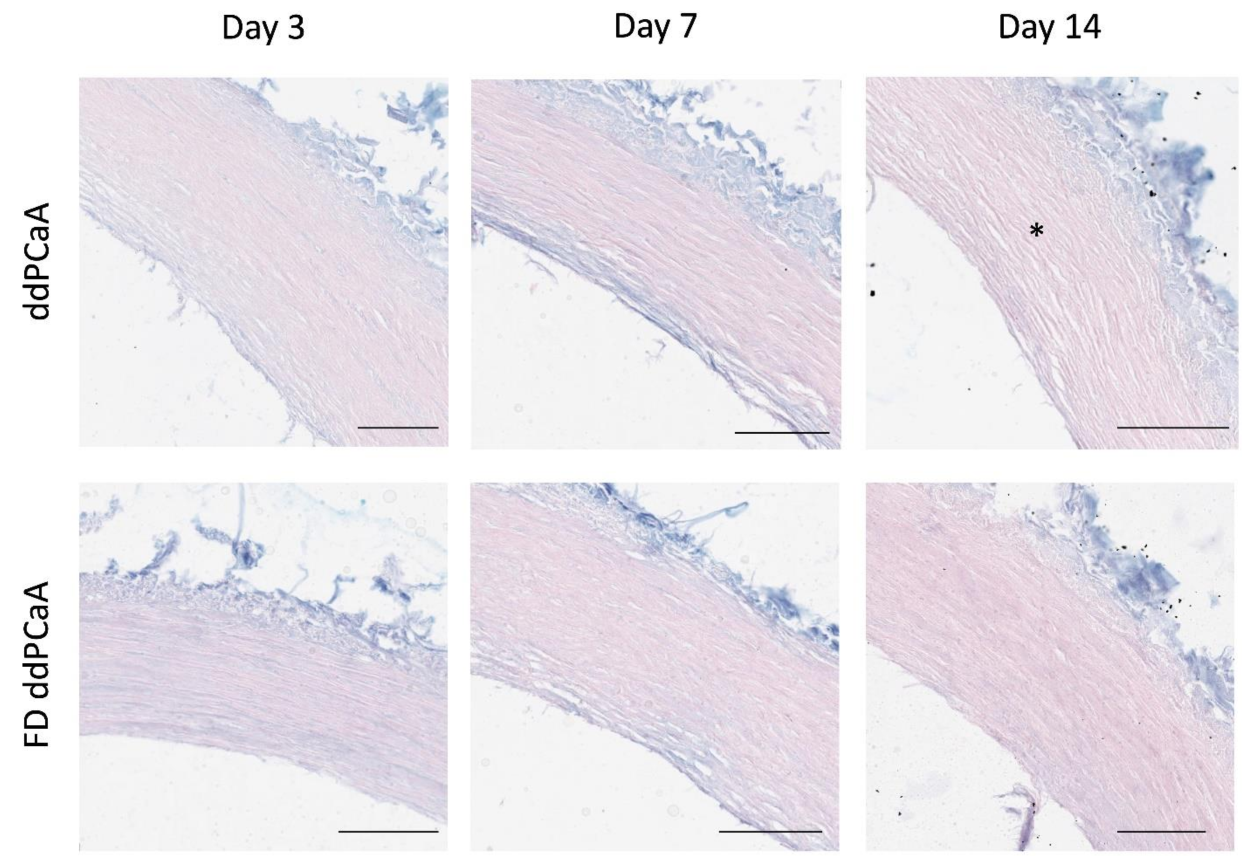

Supplementary figure 2. Alcian blue staining of unseeded vascular grafts. Scale bars are 200 µm. Asterisks marks visible intra-wall separation in comparison to the more densely packed FD ddPCaA below.
